# STIMULUS: Noninvasive Dynamic Patterns of Neurostimulation using Spatio-Temporal Interference

**DOI:** 10.1101/216622

**Authors:** Jiaming Cao, Pulkit Grover

## Abstract

Using a systematic computational and modeling framework, we provide a novel *Spatio-Temporal Interference-based stiMULation focUsing Strategy* (STIMULUS) for high spatial precision noninvasive neurostimulation deep inside the brain. To do so, we first replicate the results of the recently proposed temporal interference (TI) stimulation (which was only tested in-vivo) in a computational model based on a Hodgkin-Huxley model for neurons and a model of current dispersion in the head. Using this computational model, we obtain a nontrivial extension of the 2-electrode-pair TI proposed originally to multielectrode TI (*>* 2 electrode pairs) that yields significantly higher spatial precision. To further improve precision, we develop STIMULUS techniques for generating spatial interference patterns in conjunction with temporal interference, and demonstrate strict and significant improvements over multielectrode TI. Finally, we utilize the adaptivity that is inherent in STIMULUS to create multisite neurostimulation patterns that can be dynamically steered over time.

## 1 Introduction

High precision noninvasive sensing and stimulation of the brain can transform neuroscience, brain-machine interfaces, and clinical treatments of neural disorders. In the context of current stimulation, a limitation in stimulating a localized brain region arises from the physics of wave travel in any dispersive medium. The amplitude of currents is larger for neurons closer to the electrodes. For classical Transcranial Direct Current (DC) and Alternating Current (AC) stimulation, this means that to stimulate neurons farther away from the electrodes, one ends up stimulating closer neurons as well. Interestingly, recent work of Grossman et al. [1] shows that it is possible to engage neurons far from the electrode without stimulating those close to the electrodes. To accomplish this, they develop novel “Temporal-Interference” (TI) stimulation strategies, and validate their efficacy in mice models. This approach holds promise, but, as the authors of [1] themselves note, its precision and depth need to be dramatically improved for it to be applicable to humans and other primates.

In this paper, we first obtain a deeper understanding of TI stimulation by replicating results of [1] in a computational model. We note that this replication itself is significant: results in [1] are obtained through in-vivo mice experiments, and it is important to be able to understand them through existing computational models of neurons. Further, this understanding enables us to improve on the TI stimulation strategies. The first strategy we obtain is a generalization of TI stimulation to multiple (*>* 2) electrode pairs, which provides significantly higher spatial resolution than the 2-electrode-pair TI proposed in [1]. Nevertheless, for larger heads (and thicker skulls), the inevitable dispersion of currents with distance still reduces precision of temporal interference. To actively combat this loss in precision, we propose “Spatio-Temporal Interference-based stiMULation focUsing Strategies” (STIMULUS) that, somewhat counterintuitively, *harness* the current dispersion (or more precisely, the spatial diversity of current dispersion) to *improve* the spatial precision of stimulation. We next review the fundamentals of TI stimulation [1], examine its limitations, and describe how our strategy is capable of overcoming some of these limitations.

**Review of TI stimulation:** TI stimulation [1] is based on simultaneous application of two sinusoidal currents of slightly different frequencies to produce temporal interference patterns. In mice models, Grossman *et al*. observe that neurons near the surface are not stimulated despite stimulation of neurons deeper in the brain. The key intuition behind TI stimulation is as follows (see Fig. 3): it is well known that neurons do not respond to high frequency sinusoidal current stimulation [2] in part because of high capacitance of neural membranes. However, at locations where the sinusoidal currents have comparable amplitudes, addition of two high-frequency sinusoidal currents of slightly different frequencies produces a waveform that is a high-frequency “carrier wave” (corresponding to the average of the frequencies of the two sinusoids) modulated by a low-frequency envelope oscillating at the “beat” frequency. That is, the envelope’s frequency is the difference of the frequencies of the input sinusoids. This slow wave of the envelope is able to successfully engage neurons whose membranes may not respond to the high-frequency carrier waves. At locations where the amplitude of one sinusoid dominates the other (e.g. closer to one of the electrodes), the envelope does not oscillate significantly, and the neurons were observed to not fire.

That the engaged neurons are observed to fire at beat frequencies is significant: if the strategy was simply utilizing the low-pass filtering effect of the neural membrane [2], then it would not engage neurons at all because a low-pass filtering of a superposition of two high-frequency sinusoids would (ideally) yield a zero output. Fundamentally, the strategy must exploit a nonlinearity in the neural membrane potential in response to the input current. The authors themselves speculate that the neuron might be performing an operation similar to envelope demodulation. Interestingly, a form of nonlinearity that is relevant to *nonideal* envelope demodulation, namely, rectification (instead of ideal envelope demodulation that utilizes Hilbert transforms [3]), has been noted in the literature [4, 5]. Our observations in Section 3.1.1 suggest that indeed, it is the nonideal envelope demodulation that the neurons seem to be performing because the demodulation is affected by the center frequency.

**Limitations of the strategy employed in [1]:** (i) To not stimulate neurons along the plane equidistant from the two electrode pairs (see the *x* = 0 plane in Fig. 8 (a), “TI: 2 pairs”), the TI-stimulation technique in [1] depends solely on current dispersion in the medium and placement of the electrodes. Along the *z*-axis in Fig. 8 (a), the strategy requires currents to concentrate quickly as they get shallower. However, for thick skulls, current disperses significantly as it leaves the skull and enters the brain^1^, and thus currents are not very concentrated in shallow regions, reducing the spatial precision substantially. Along the *y*-axis, where the amplitudes of the two sinusoids remain equal, TI stimulation requires the current density to decay quickly to limit the region of stimulation. Again, for media in which current has dispersed significantly before entering the brain, current density will not decay quickly as one travels away from the origin along *y*-axis. (ii) For larger heads, an additional difficulty arises: because of increased current dispersion, supplied currents also need to be large to ensure that desired neurons at depth are engaged. Not only can large currents test limits of tissue damage, prior work [2] as well as our observations on the HH model (Section 3.1.1) indicate that this could lead to engagement of neurons close to the electrodes where the current densities are higher.

This work addresses these critical limitations of the technique in [1], and also develops techniques for *multisite* and *steerable* stimulation.

**Our contributions:** Intuitively, the spatial-interference aspect of our technique is used to limit the spatial extent of stimulation along the *z*-axis, while we advance on the temporal-interference aspect to reduce stimulation along the *x*- and *y*-axes. We now describe steps that help us accomplish this increase in spatial precision:

i. **Modeling and replicating results of [1] in a simulated model**: To accomplish this, we use the Hodgkin-Huxley (HH) model [8] and simplified models of spatial dispersion of currents in tissue. These results show that TI-stimulation results of [1] can be replicated using these simplified models, which opens the door to use of computational strategies in examining and improving TI-stimulation.
ii. **Multielectrode TI stimulation**. Using the HH model, we observe that very low frequency TI envelope modulations require higher current densities for stimulation (see Fig. 4). This observation leads us to propose novel multi-electrode-pair TI strategies that attain higher precision stimulation than the the 2-electrode-pair strategies in [1]. Our strategies involve having small differences in frequencies in nearby electrode-pairs, and large differences in frequencies in farther electrode-pairs. We show in Section 3.1.2 that this ensures more focused stimulation in the *z*=0-plane than the original 2-electrode TI stimulation.
iii. **STIMULUS**: STIMULUS goes a step further and harnesses spatial diversity to generate “current lenses” that focus a particular sinusoidal current at desired points. One can think of this strategy as replacing an electrode-pair in multielectrode TI with a “patch” of multiple electrode-pairs, with each electrode-pair in each patch generating currents of the same frequency. These patches act as current lenses, and the combined effect of these lenses improves resolution of multielectrode TI. We generate spatial interference patterns by exploiting knowledge of decay of currents in the brain tissue to focus the signal of each patch near the desired site of stimulation. This is an adaptation of the concept of “beamforming” in spatial filtering [9, 10, 11] to the case where multiple beamforming patches are used. We combine this with multielectrode TI to create stimulating temporal interference patterns the focus region.

Section 4 shows that STIMULUS also obtains dynamic steerability of stimulation (without moving the electrodes themselves), and can obtain simultaneous stimulation at multiple sites providing rich patterns of stimulation. Steerable multisite stimulation is important in many applications, e.g. in providing feedback in brain-machine interfaces, and in peripheral nerve stimulations [12].

Conventionally, electrodes used in transcranial Direct Current and Alternating Current Stimulation (tDCS and tACS, respectively) are arranged as rectangular patches, which tend to have poor focusing precision [13, 14]. Recent works have shown that carefully chosen electrode montages are capable of generating more focused stimulation (e.g. [13, 15]). Another class of strategies use a larger number of electrodes (8 in [10], 64 in [11], and as many as 336 in [16]) whose locations are guided by EEG electrode locations. They then solve optimization problems to tailor stimulation patterns. However, unlike results in this paper, these approaches [10, 11, 16] utilize only spatial interference, and are unable to get high precision while stimulating deep sites.

One limitation of our work is that we consider idealized spherical head models. However, given the success of beamforming-based current stimulation in real-head models [10, 11], we expect to be able to generalize these techniques to real heads. Other shortcomings of our work, and ongoing as well as future work directions, are discussed in Section 5.

## 2 Model, methods and notation

We use a 3-sphere head model [6]. We denote the brain’s conductivity by *σ*_1_, the skull’s by *σ*_2_, and the scalp’s by *σ*_3_. We largely assume that *σ*_1_ = *σ*_3_. The ratio *σ*_2_*/σ*_1_ is assumed to be either 1/80 [6] or 1/15 [17], which span the range of skull conductivities noted in the existing literature. The current dispersion is modeled through a finite-element model that incorporates conductances in each layer, and boundary conditions are provided by input currents from the electrodes. We assume a purely resistive medium for current dispersion, as has been observed in, e.g., [18]. The model is illustrated through examples in Fig. 1.

**Figure 1:**
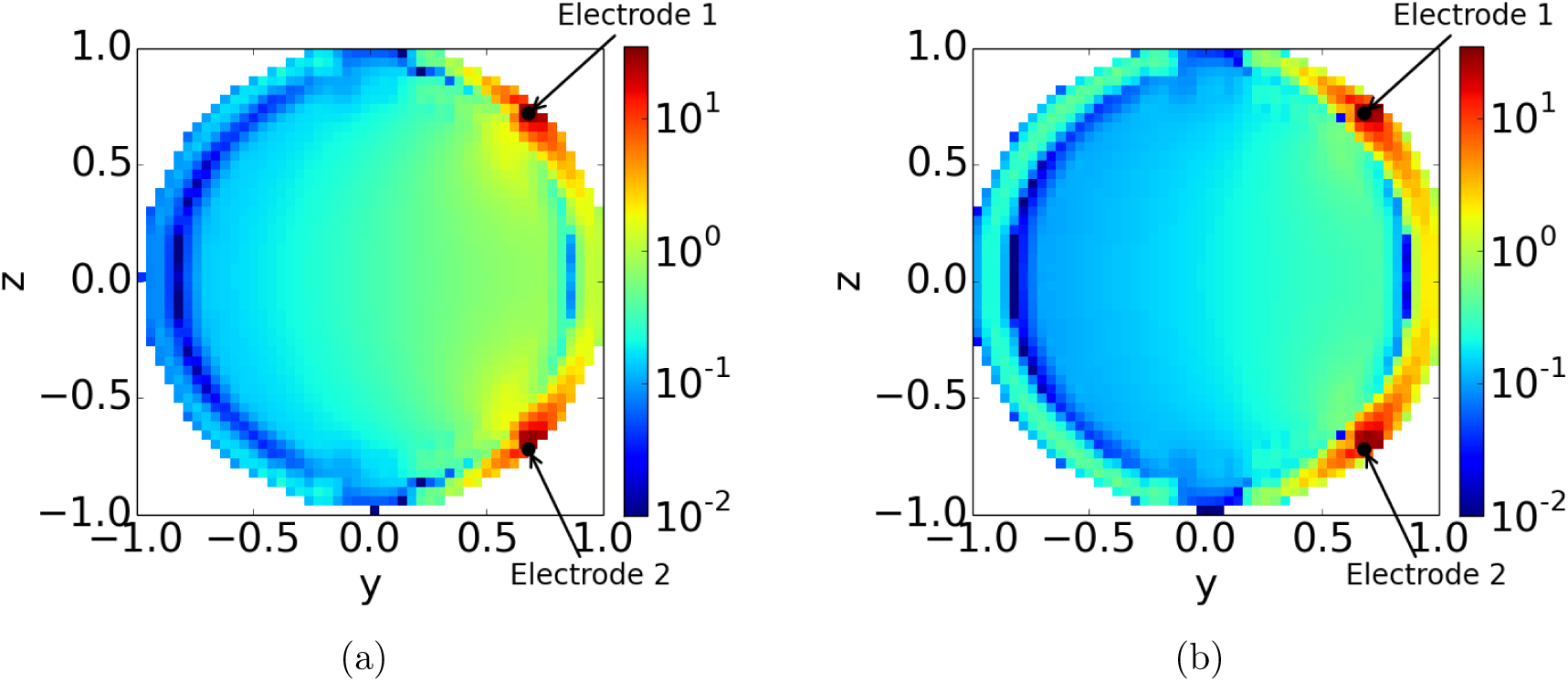
The figure shows the spatial diversity in current density (in *µA* per unit area) for a total current of 1*µA* from the electrodes in a unit-sphere head with the radii of the brain, skull, and scalp spheres being 0.8, 0.9, and 1.0 unit. Electrodes are placed at *x* = 0*, z* = *±*0.72 on the outermost sphere of radius 1, which corresponds to polar angles of *θ ≈ π/*4 and 3*π/*4. The slice shown is along the plane *x* = 0. (a) assumes that *σ*_2_*/σ*_1_ = 1/15. (b) assumes that *σ*_2_*/σ*_1_ = 1/80. A visual examination of the figures reveals the spatial diversity of currents, as well as fact that within the brain (e.g. see along the *z* = 0 plane), there is higher dispersion with *σ*_2_*/σ*_1_ = 1/80.

The electrode pairs are assumed to be placed so that the line joining the electrodes is parallel to polar axis. The current density map for a single pair of electrodes in this model is shown in Fig. 1. For multiple electrode pairs, the current density is simply a sum of current densities due to individual electrode pairs. For sinusoids of the same frequency (say *f*), this summation simply results in a sinusoid of frequency *f* at every location in the brain, with amplitude and phase that depend on the location of the point vis-a-vis that of all of the electrodes.

We use the well known Hodgkin-Huxley (HH) model [8] for understanding stimulation under current inputs. For simplicity, we assume that we know the orientation of the neuron to be stimulated, and that all neurons in the brain are described by the same Hodgkin-Huxley parameters and the same orientation^2^. More work, including allowing for different neural orientations, is needed to fully understand the accuracy of our strategies.

We note that when keeping input current (both amplitude and electrode-pair angular location) constant, changing radius of the head (ratios of radii of three layers remaining the same) is equivalent to simply multiplying the current densities inside of the head by the same scaling constant. This is because if we keep the topology of the finite element model the same, current in any discrete voxel for a given input current does not change. What changes is the unit area, which is scaled according to voxel size. For the rest of this paper, we choose the head to have a radius of 10 cm, which is close to that of human head [6]. Therefore, 1 unit length in figures as well as coordinates correspond to 10 cm, and this is used when converting current densities to total input currents.

*Methods*: For each simulation, we solve Hodgkin-Huxley equations using Dormand-Prince method [20] for a total span of 1000ms sampled at 25 kHz.

In order to estimate firing, we apply a sixth-order Butterworth bandpass filter to keep only frequency components between 100 and 1000Hz. We adopt a simple criterion to decide firing: when the filtered membrane potential crosses a threshold of 30mV, we deem it as a firing event. To avoid boundary effects, we removed the first and last parts of potential time trace. The 30mV threshold is empirically observed to obtain “robust” firing for our HH model. There are cases when membrane potentials show a “tendency” to fire by showing small peaks in the filtered response, yet the peaks are not pronounced enough for us to classify them as firing (e.g. see Fig. 3b and Fig. 6b).

To demonstrate our improvements, we assume for most of our initial (single, fixed focus) results that the stimulation is required at the deepest point: the center of the spherical head model. Further, we impose the constraint that the stimulation pattern include a “disk” of radius approximately 2 cm at the head center. This is motivated by the observation that in deep-brain stimulation, one may be interested in stimulating only a deep region in the brain, (e.g. the hippocampus, but not the tail of caudate nucleus which is just above the hippocampus). More importantly, it forces all stimulation strategies to stimulate a minimal region, not just a single point. The latter can yield misleading results because strategies can tailor current waveforms to barely stimulate one point in the space, but in practice, small uncertainty in conductivity parameters and neural location can render these strategies ineffective.

In the remaining sections, we use both Cartesian coordinate system (*x, y, z*), and polar coordinates (*r, θ, φ*). We use bold font to denote vectors, and uppercase boldfont to denote matrices. For any matrix **A**, **A**^*T*^ denotes its transpose.

## 3 Improving spatial resolution of noninvasive neurostimulation for a single deep focus

In this section, we aim to obtain strategies that harness spatial diversity of current dispersion to generate spatiotemporal patterns of current stimulation that improve resolution of noninvasive neurostimulation.

### 3.1 Multielectrode TI stimulation

We first obtain a multielectrode extension of TI stimulation. It turns out that this extension is nontrivial: it requires understanding envelope waveforms and carrier frequencies that lead to stimulation, as well as those that do not. This understanding, combined with an understanding of spatial dispersion of currents with distance, leads us to multielectrode TI strategies. First, however, we show that the 2-electrode-pair TI stimulation obtained in mice models in [1] can be replicated in a Hodgkin-Huxley-model based simulator.

#### 3.1.1 Understanding the 2-electrode-pair TI stimulation using a Hodgkin-Huxley model-based simulator

We first analyze the HH model to understand how neurons respond to slow envelopes on high-frequency carrier waves. To do so, we use the Hodgkin-Huxley model with parameters from [21].

**Low-carrier frequencies**: We observe that TI stimulation is not effective at low carrier frequencies, e.g. 300 Hz or even up to 1500 Hz, for these HH model parameters. While the interfering pattern can be made to stimulate at the sphere center, we observe excessive undesired stimulation. In fact, neurons at all points connecting the sphere center and the electrodes appear to be stimulated, as shown in Fig. 2. We believe this is because the carrier frequency is too low, and the neurons can still follow it. Thus, the neurons are not responding to just the envelope.

**Figure 2:**
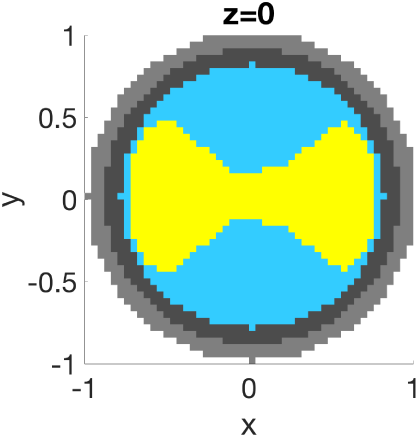
Temporal interference does not work when frequencies of base sinusoids are not very large. In this example, frequencies are chosen to be 290 Hz and 300 Hz, and they both generate current density of 10.6*µA/cm*^2^ at the center, the minimal required intensity for robust stimulation at the center. It turns out that moving away from center, closer to the electrodes, does not reduce firing.

**High-carrier frequencies**: However, at carrier frequencies larger than 1800 Hz, we observed that TI stimulation is effective in that moving away from the site of equal amplitude of sinusoids causes the neurons to stop firing. This is desirable for being able to stimulate deep within the brain without shallow stimulation. This is shown in Fig. 3 for two electrode pairs which generate currents of frequencies 1990 Hz and 2000 Hz.

**Figure 3:**
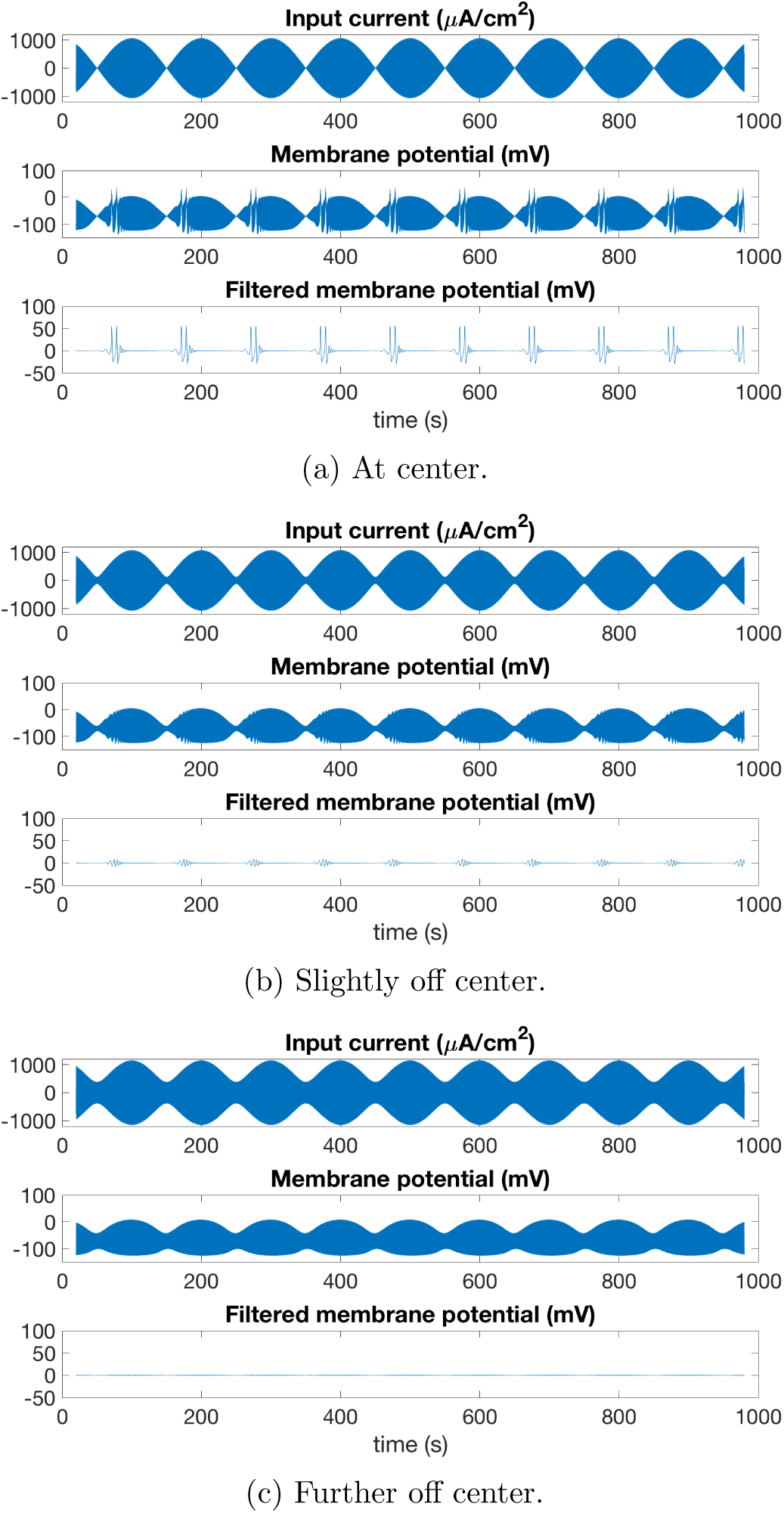
Understanding two-electrode-pair TI stimulation for *σ*_2_*/σ*_1_ = 1/80. The figure shows input current, membrane potentials, and filtered membrane potentials (filtered to elicit firing) of neurons at different locations. (a) shows results at the sphere center, where both sinusoidal currents have equal amplitudes. The envelope of input current in this case has maximum modulation depth as it touches 0. Clear evidence of neuron firing can be observed in both the membrane potential and the filtered membrane potential. (b) shows results at a point slightly away from center ([0.12, 0, 0]), where amplitudes of two sinusoids are slightly different such that the envelope of the input current does not fall to zero. Membrane potential both before and after filtering shows that the neuron is close to firing, yet is unable to. (c) Shows results at a point farther away from center ([0.36,0,0]), where two sinusoids have substantially different amplitudes, causing the envelope to be rather flat and far from zero. The neuron does not show any evidence or tendency of firing in this case.

We note that *this change in firing behavior as a function of carrier frequency suggests that the membrane potential is not well approximated as ideal envelope demodulation*. It appears that the simpler diode-resistor-capacitor model for a nonideal envelope detector [3] might be a better approximation, instead of the ideal Hilbert transform envelope detector. Nevertheless, the precise impact of nonlinearity of neural membrane’s potential as a function of the input current remains to be fully understood, and it is plausible that waveforms other than interfering sinusoids might be more effective at exploiting this nonlinearity.

We now explain in some detail how we are using the Hodgkin-Huxley model to infer locations of neural stimulation. At the sphere center, where both sinusoidal currents have equal amplitude (Fig. 3a), robust firing can be achieved, as can be inferred from bandpass filtered membrane potential [1]. If we probe a neuron far from center (e.g. moving along the *x*-axis in Fig. 3c), the currents have unequal amplitudes and the envelope has less “depth” in its “valleys”. The filtered membrane potential shows that the firing stops in this case. This observation fully agrees with those reported in [1]. For the chosen location in Fig. 3b, the probed neuron lies in between the above two. Here, the envelope has a larger (but not large enough) difference between peaks and valleys, and from the filtered membrane potential, it appears as if the neuron is close to firing, but does not fire. The membrane potential has some high frequency components, but the amplitudes are small compared to Fig. 3a.

The resulting stimulation is shown in Fig. 7 for *σ*_2_*/σ*_1_ = 1/80 and Fig. 8 for *σ*_2_*/σ*_1_ = 1/15 along the planes *x* = 0, *y* = 0, and *z* = 0. For any one pair, one electrode is placed at *θ* = *π/*4 and the other at *θ* = 3*π/*4 on the *y* = 0 plane. Observe that for less dispersive skulls, undesirable (albeit limited) shallow current stimulation is observed in conjunction with deeper stimulation.

#### 3.1.2 Multielectrode TI stimulation

Let us first focus on just two interfering sinusoids as the input current to the neuron, one at 2000 Hz and the other at (2000 + ∆*f*) Hz for varying ∆*f*. In this case, Fig. 4 shows that the threshold current density needed to achieve effective TI stimulation is higher when the difference in frequency is smaller. It seems to us that for this input current, the neural membrane’s response function can be interpreted as an envelope-detector followed by a band-pass filter. For this configuration, the smallest required current density is at ∆*f* = 30 *−* 40 Hz, and hence the neurons are easiest to stimulate at that frequency difference. However, more importantly, observe that the curve has a steep rise as the frequency is lowered below 10 Hz.

**Figure 4:**
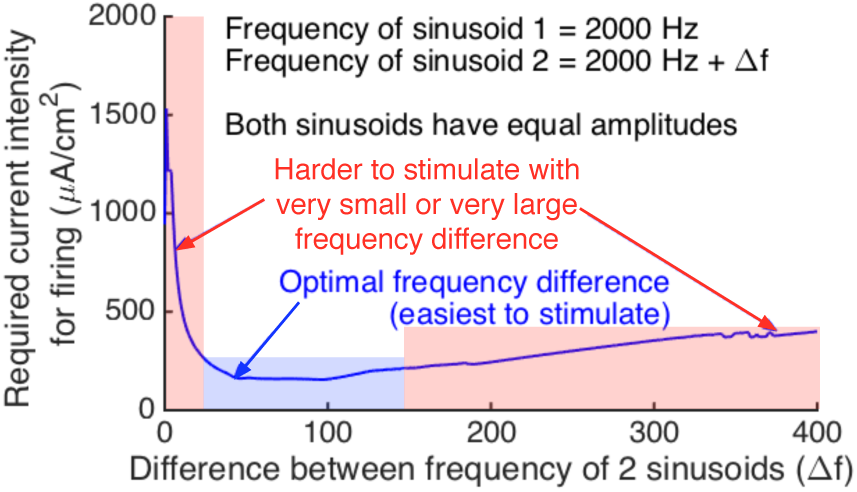
Minimum current density of envelope-modulated signal needed for 2-electrode TI stimulation for a Hodgkin-Huxley neuron with model parameters drawn from [21]. The figure shows that there is an optimal frequency difference of the interfering sinusoids (both of equal amplitude) to enable stimulation at low current densities. We utilize observation in the design of our multielectrode TI strategy.

This observation suggests an interesting strategy for frequency allocation and electrode-pair placement for multielectrode TI stimulation. First, for electrode-pair placement, we simply place electrode-pairs as shown in Fig. 5 where each electrode pair has exactly one electrode in the top hemisphere, and the electrodes in the top hemisphere are arranged in a ring. Electrodes in the bottom hemisphere and their top hemisphere counterparts are symmetric about *z* = 0 plane. This electrode pair arrangement can generate higher current densities at the sphere-center.

**Figure 5:**
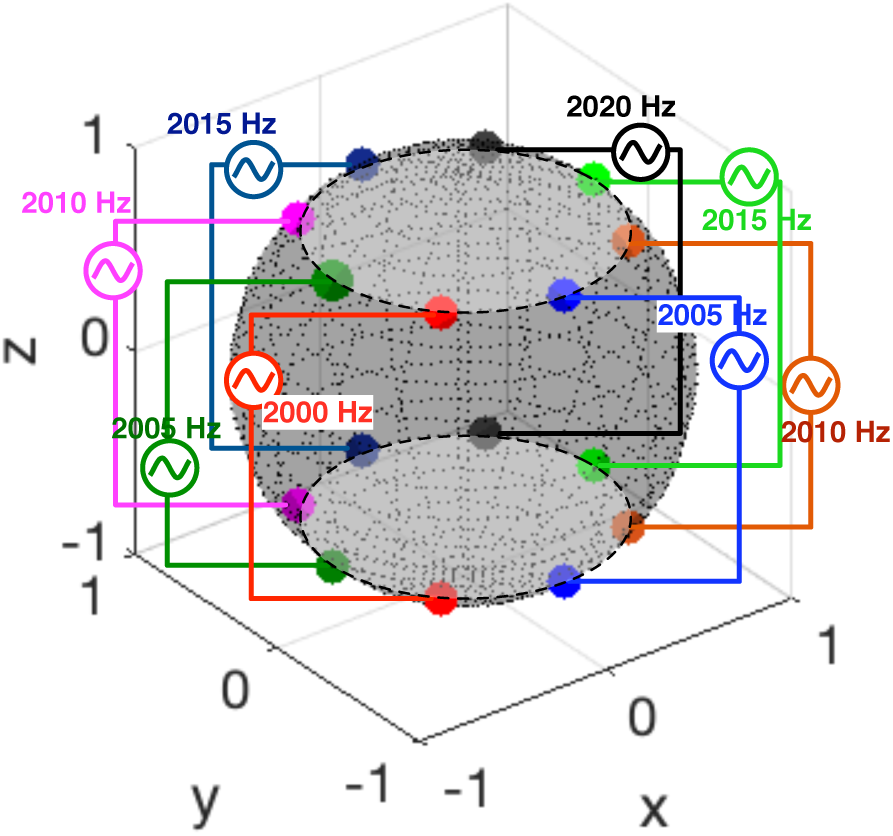
Example electrode-pair placement and frequency allocation for multi-electrode-pair TI for stimulating at the sphere center. Example frequency allocation is also shown. Nearby electrode-pairs frequencies are close to each other so that neurons closer to the electrode pairs from the sphere center are not stimulated.

We now discuss frequency allocation for the electrode-pair placement discussed above. The goal is to have the neurons away from the center to not fire while the neurons at the center do. If one allocates frequencies so that nearby electrode-pairs have frequencies that are close to each other (e.g. 1-5 Hz), but diametrically opposite electrode-pairs can have large differences (e.g. up to 20 Hz), then away from the center, the dominant sinusoids have frequencies close to each other, lowering the higher-frequency-content (10 *−* 20 Hz) of the envelope, which makes firing less likely to happen. However, close to the center, because all sinusoids are adding up with approximately equal amplitudes, the envelope can have significant high frequency content. Analogous to 2-electrode-pair TI (Fig. 3), we illustrate Hodgkin-Huxley neuron responses at different spatial locations in Fig. 6. Here, we make similar observations for a more complex envelope: stimulation is more likely to happen when the envelope is less flat.

**Figure 6:**
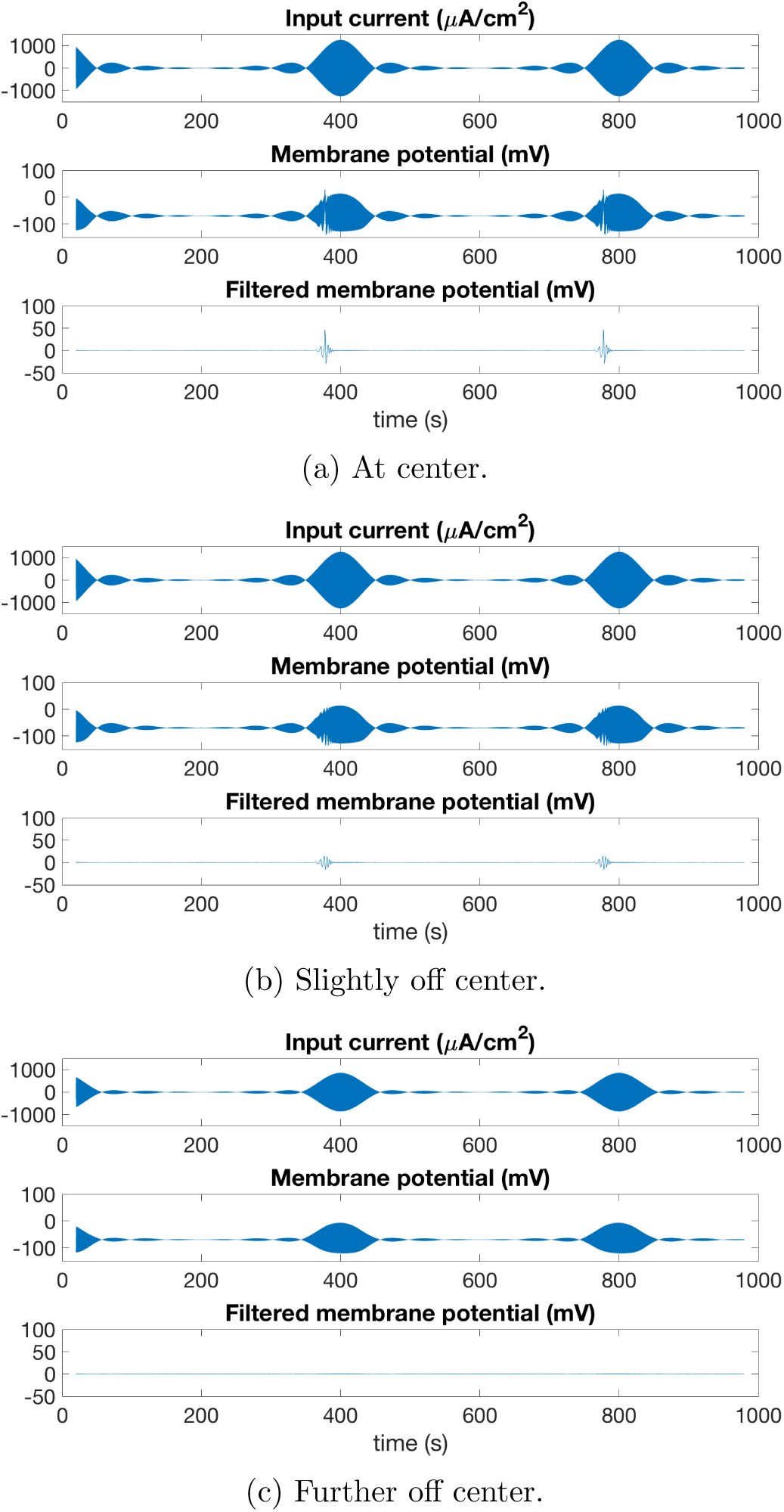
Understanding sixteen-electrode-pair TI stimulation for *σ*_2_*/σ*_1_ = 1/80. The figure shows input current, membrane potentials, and filtered membrane potentials (filtered to elicit firing) of neurons at different locations. (a) shows results at the spherical center, where all sine waves have equal amplitudes. Envelope in this case has 0 DC bias. Clear evidence of neuron firing can be observed. (b) shows results at a point slightly away from center ([0.16,0,0]), where amplitudes of sinusoids are slightly different such that there is a small amount of DC value in summed input. Membrane potentials show that the neuron has a tendency to fire, yet the effect is not evident enough. (c) shows results at a point farther away from center ([0.76,0,0]), where sinusoids have very different amplitudes and lead to large DC component in summation. The neuron does not show any evidence or tendency of firing in this case.

Indeed, this strategy succeeds in outperforming 2-electrode-pair TI with improved precision of stimulation in the *z* = 0-plane, as shown in Fig. 7 for *σ*_2_*/σ*_1_ = 1/80, and Fig. 8 for *σ*_2_*/σ*_1_ = 1/15.

**Figure 7:**
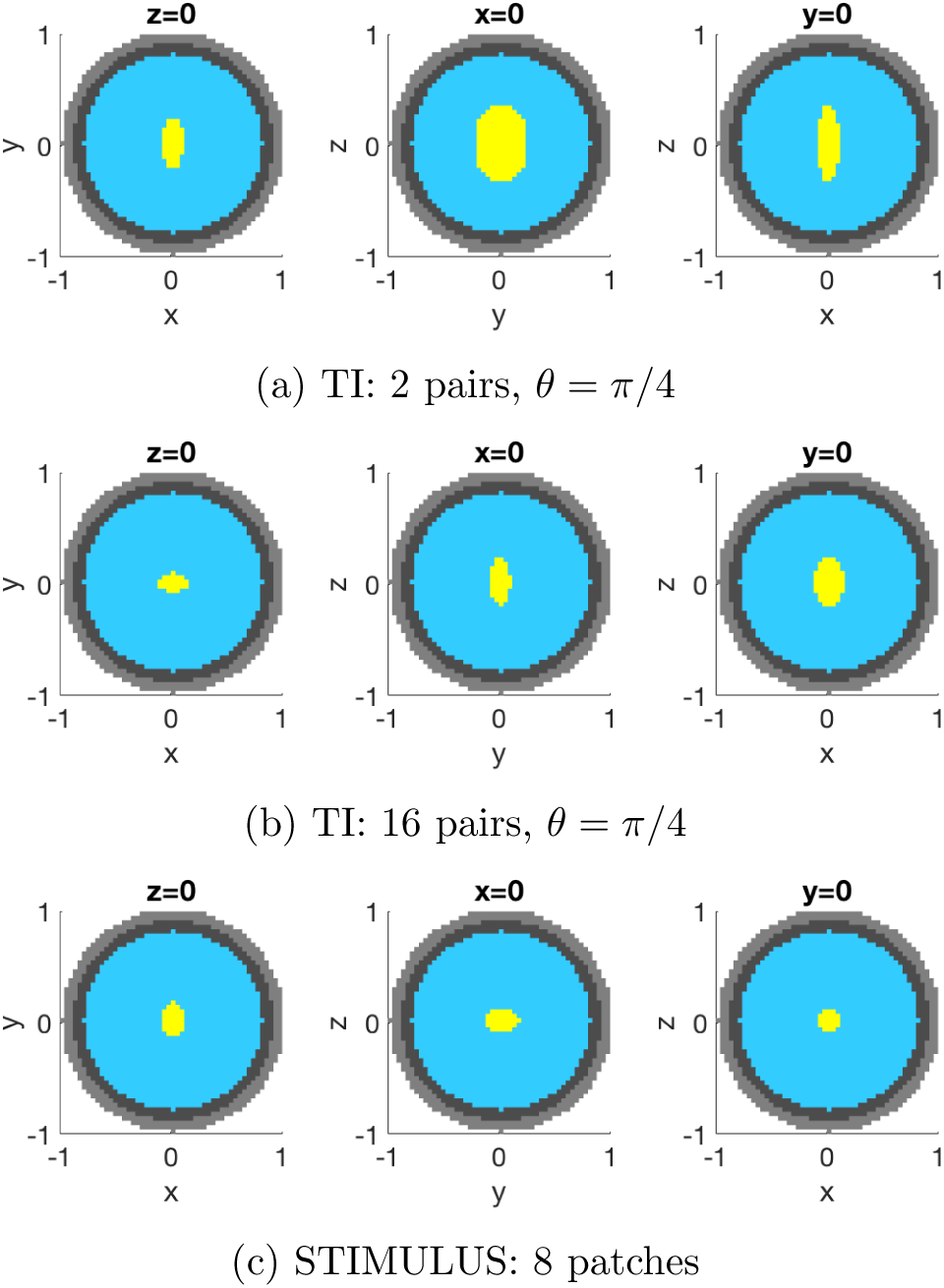
Stimulation regions using TI, multielectrode TI, and STIMULUS, assuming *σ*_2_*/σ*_1_ = 1/80. (a) TI stimulation using 2 pairs of electrodes in *y* = 0 plane does not provide good precision along the *z*-axis; (b) Multielectrode TI using 16 pairs of electrodes provides improved focus, but the focus along the *z*-axis is still poor; (c) STIMULUS using 8 patches reduces region of stimulation along the *z*-axis by Blue: no stimulation; Yellow: stimulation; Dark gray: skull; Light gray: scalp.

**Figure 8:**
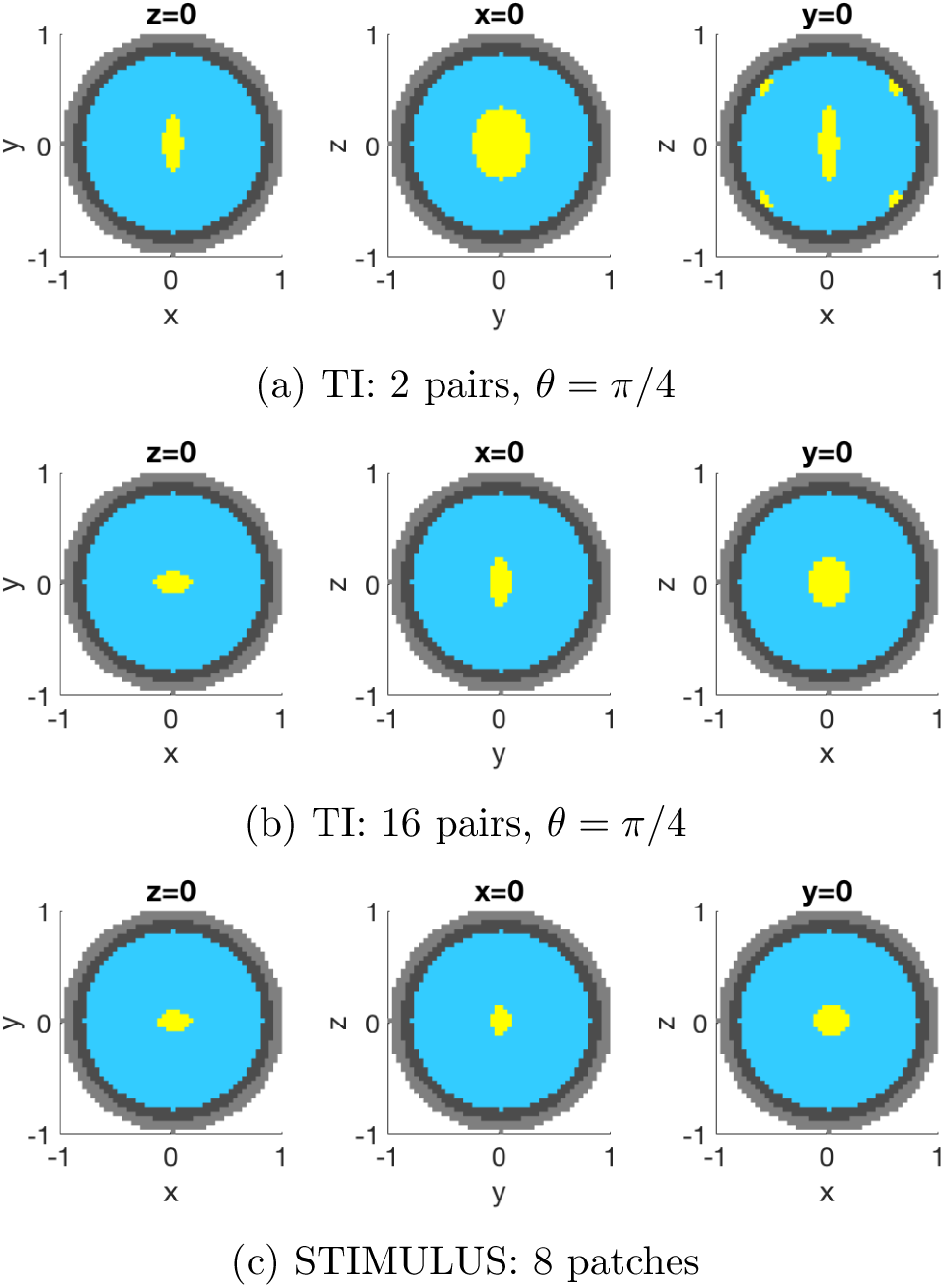
Stimulation regions using TI, multielectrode TI, and STIMULUS, for a higher skull conductivity, where *σ*_2_*/σ*_1_ = 1/15 (as compared to Fig. 7, where *σ*_2_*/σ*_1_ = 1/80). This leads to less current dispersion, and increased diversity of CID-factors. Therefore, SIMULUS, which harnesses spatial diversity, now outperforms multielectrode TI by an even larger factor. Also note the shal-low stimulation associated with 2-electrode-pair TI, which is eliminated by multielectrode TI and STIMULUS. Blue: no stimulation; Yellow: stimulation; Dark gray: skull; Light gray: scalp.

However, the attained precision along the *z*-axis is still not very high because this strategy does not actively limit the spread of stimulation region in that direction. It is possible that further increase in number of electrodes using multi-TI stimulation could improve precision along the *z*-axis, but, extrapolating from Fig. 7 and Fig. 8, it would come at the cost of reduced firing rate. We now describe how SIMULUS exploits spatial diversity and combines beamforming approaches with multielectrode TI to limit spread along the *z*-axis.

### 3.2 Harnessing spatial diversity using STIMULUS

In the electrode-pair configurations in the multielectrode TI of Section 3.1.2, the extent of stimulation along the *z*-axis (i.e., the polar axis) depends only on electrode placement. The strategy relies on the physics of the problem (i.e., concentration of currents near the the electrodes, see Fig. 1) to ensure that there is sufficient decay to limit stimulation extent along the *z*-axis. This reduces the precision along the *z*-axis, and the reduced precision could be unacceptably low for applications such as deep-brain stimulation.

Here, we propose our alternative strategy – STIMULUS – that generates spatio-temporal patterns of interference to improve resolution along the *z*-axis. One can think of this strategy as replacing an electrode-pair in multielectrode TI with a “patch” of multiple electrode-pairs, with each electrode-pair in each patch generating currents of the same frequency. It is instructive to think of each patch of electrodes as a “computational current lens,” that attempts to focus currents (using constructive interference) at some desired points of focus, and cancel currents (using destructive interference) at a few other (carefully chosen) points.

We now formally describe how these constraints of constructive and destructive interference can be written down in an optimization formulation. Assume that we have *p* electrode-pairs in each patch with current-amplitudes **x** = [*x*_1_*, x*_2_*, …, x*_*p*_]^*T*^. Let us assume that there are *n* focus points for this patch, and *m* points where we minimize current densities to suppress stimulation.

Let us define **current intensity decay (CID) factor** as the ratio of the generated current density at a point to the total input current from an electrode pair that generates that current density. Thus, this factor depends both on the location of the point at which the current density (due to just the chosen electrode pair) is being estimated, and the location of the electrode pair generating currents^3^. Let **A**_focus_ and **A**_cancel_ denote the CID-factor matrices, formed by the focus and cancellation points respectively, where **A**_focus_ is a *p × m* matrix and **A**_cancel_ is a *p × n* matrix with each column being a vector containing the CID factors from all electrodes to a focus point (for **A**_focus_) or a cancel point (for **A**_focus_). Therefore, the total current intensity at focus points can be written as 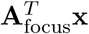, and at cancel points, 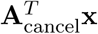.

For computational tractability, we formulate our objective function as minimization of the square sum of currents at all cancel points, 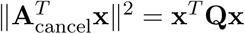 where 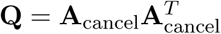.

For simplicity, we constrain the optimization problem to have each patch produce the same current density *I*_*stim*_ at all focus points. The value of *I*_*stim*_ for optimization is the required current density obtained through single-neuron Hodgkin-Huxley simulations. More precisely, for an input that is a sum of sinusoidal current densities (each of value *I*_*stim*_), with each sinusoid’s frequency corresponding to the frequency of a patch in the system, we find the minimum *I*_*stim*_ that ensures robust stimulation in the region of interest.

This leads us to the following optimization problem that yields the optimal current allocation **x**^*∗*^ at each electrode-pair in the patch:

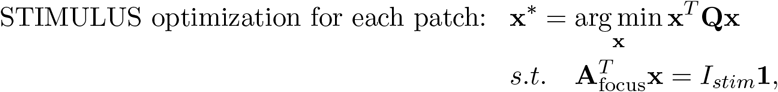

where **1** is the all-ones vector.

In practice, one might need to further constrain the problem, e.g., by limiting currents from each electrode-pair, and further simplify the problem by recognizing that approximate cancellation is still helpful.

We demonstrate improvement of precision using STIMULUS for different values of skull conductivity. The resulting focus regions for very dispersive skull (*σ*_2_*/σ*_1_ = 1/80) and a less dispersive (*σ*_2_*/σ*_1_ = 1/15) are shown in Fig. 7 and Fig. 8 respectively. In both cases, we observe that the stimulation along *z*-axis is more focused in comparison with multielectrode TI. In order to obtain these results, centers of patches are placed at regular *φ* intervals, and in polar direction they all span from poles (*θ* = 0) to (*θ* = 5*π/*12, i.e., *π/*12 away from equator) in the top hemisphere, with symmetric placement (about *z* = 0 plane) in the bottom hemisphere. For each patch, *p* = 100 locations in the top hemisphere are chosen randomly as electrode locations.

For *σ*_2_*/σ*_1_ = 1/80, we let the patch at *ϕ* = 0 focus at [*−*0.2, 0, 0], the patch at *ϕ* = *π/*2 focus at [0, 0.08, 0], the patch at *ϕ* = *π* focus at [0.16, 0, 0], the patch at *ϕ* = 3*π/*2 focus at [0*, −*0.08, 0], and others focus at the center. All patches minimize current density (“cancel”) at 0.15 above their foci. Further, all patches have *I*_*stim*_ = 199*µA/cm*^2^, with constraint that current amplitude each electrode pair should not exceed 2.985 *µA*. For focusing points, notice that some of the patches are not focusing at *x* = 0 or *y* = 0. This helps improve the precision because having all patches focus at the center leads to a larger and asymmetric pattern of stimulation. We believe that this asymmetry is due to our choice of frequency allocation, and there is room for further optimization of this strategy. But moving foci slightly farther away from the center reduces this asymmetry and also improves the precision.

Similary, for *σ*_2_*/σ*_1_ = 1/15, we let the patch at *ϕ* = 0 focus at [*−*0.16, 0, 0], the patch at *ϕ* = *π* focus at [0.16, 0, 0], and all others focus at sphere center. All patches cancel at 0.15 units above their foci, and have same *I*_*stim*_ = 181*µA/cm*^2^ with amplitude of each electrode pair being no larger than 2.715 *µA*.

Finally, we note that there is a tradeoff between number of patches and precision. If fewer patches are used, each patch can have a larger spatial extent, creating more spatial diversity for its electrodes and thus attaining better focusing and cancellation and richer patterns. But this limits the number of sinusoids being used, and thus reduces the use of multielectrode TI ideas. Thus, a balance needs to be maintained and an optimal number of patches needs to be chosen.

## 4 Steerable and multisite stimulation

**Steerable stimulation:** To demonstrate steerability using STIMULUS, it suffices to show that with the same electrode locations as above, we can stimulate areas that are not at the sphere center. To do so, in our optimization problem, we let all patches focus at [0, 0, 0.5] and cancel at both [0, 0, 0.3] and [0, 0, 0.7]. Current density of each patch at focus points is chosen to be *I*_*stim*_ = 165 *µA/cm*^2^, and the current amplitude of each electrode pair is constrained to be no more than 3.31 *µA*. Results are shown in Fig. 9.

**Figure 9:**
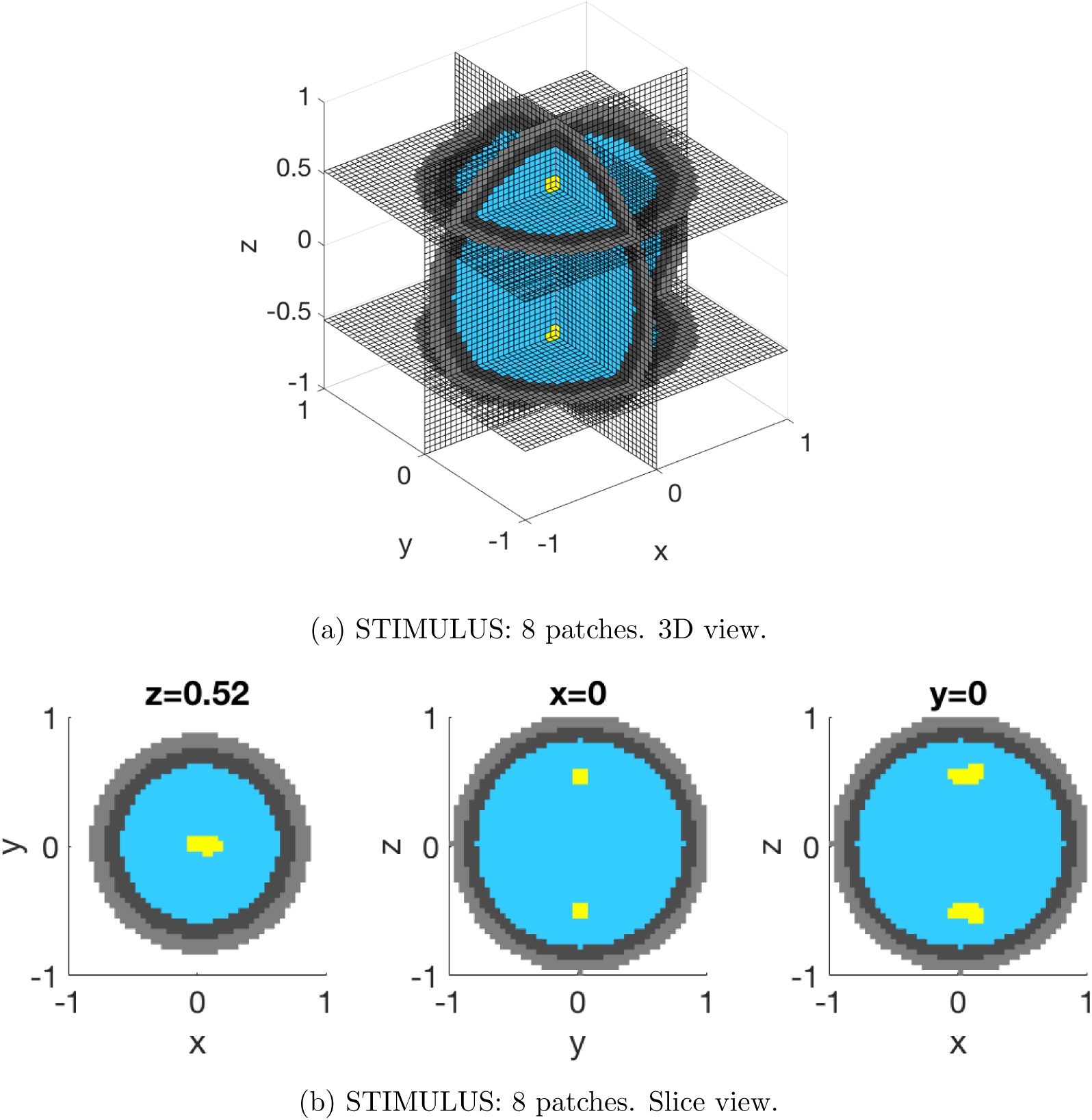
Steerable stimulation: Demonstration of 8-patch STIMULUS focusing at [0, 0, 0.5]. Notice that the slice at *z* = 0.52 is smaller because section of the sphere in that plane has a smaller radius. *σ*_2_*/σ*_1_ = 1/15, and electrode placement is the same as in Fig. 8 and Fig. 10. Stimulation is created away from center, and located only near [0, 0*, −*0.5], the focus. Mirror image of the focus, [0, 0*, −*0.5] is also observed to be stimulated, because of symmetry of electrode placement. (a) shows 3D view of stimulation pattern. Notice that the sphere center is not stimulated; (b) shows slice views of stimulation pattern, with the *z*-slice at *z* = 0.52. Blue: no stimulation; Yellow: stimulation; Dark gray: skull; Light gray: scalp.

**Multisite steerable stimulation:** We now show that STIMULUS is able to stimulate two distant sites simultaneously without stimulating regions that connect the two. We still use the same placement of electrode as for steerable stimulation above, demonstrating that STIMULUS can perform steerable multisite stimulation. We choose a shallow site, and a deep site, with all patches focusing at both [0, 0, 0] and [0, 0, 0.7], and canceling at [0, 0, 0.25] to avoid stimulating in between the two sites. We let *I*_*stim*_ of all patches be 166 *µA/cm*^2^, and the maximum allowed current of each electrode pair be 2.49 *µA*. Results are shown in Fig. 10

**Figure 10:**
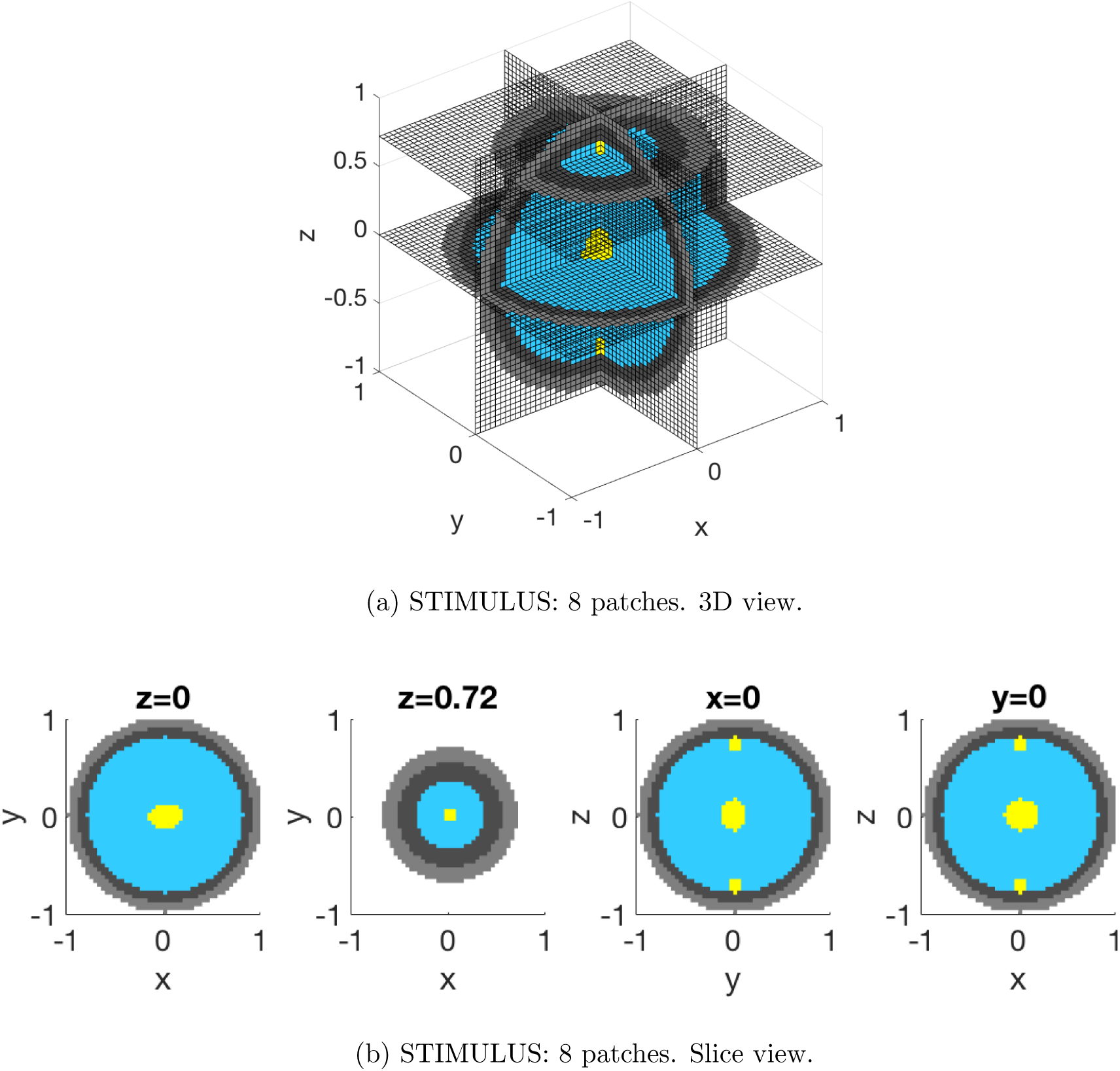
Multi-foci stimulation: Demonstration of 8-patch STIMULUS focusing at [0, 0, 0] and [0, 0, 0.7] simultaneously. *σ*_2_*/σ*_1_ = 1/15, and electrode placement is the same as in Fig. 8 and Fig. 9. Notice that the slice at *z* = 0.72 is smaller because section of the sphere in that plane has a smaller radius. Stimulation happens at center and [0, 0, 0.7] (and also [0,0,-0.7] due to symmetry of electrode placement), but neurons in between the foci are not engaged. (a) shows 3D view of stimulation pattern; (b) shows slice views of stimulation pattern, with *z*-slices at *z* = 0 and 0.72. Blue: no stimulation; Yellow: stimulation; Dark gray: skull; Light gray: scalp.

## 5 Discussion and conclusions

The novel STIMULUS strategy provided here creates patterns of spatiotemporal interference in order to improve precision of noninvasive neurostimulation. We also advance on the understanding of temporal interference stimulation, showing that it can be obtained in Hodgkin-Huxley models, and extending it to multiple electrodes. We also observe that TI stimulation relies on a nonlinearity of neuronal membrane as a function of the input current density. Thus, it cannot simply be modeled as a low-pass filter. Further, it cannot simply be modeled as an envelope demodulator either, as shown in our results for carrier frequencies that are not very high. Thus, the behavior of neural membrane as a function of input current deserves further study.

It is shown that STIMULUS outperforms multielectrode TI in its precision, and also eliminates shallow firing. Multielectrode TI has two shortcomings. First, despite that multielectrode TI is capable of reducing undesired firing, we notice that improvement in spatial precision is limited. Second, although using larger number of electrodes can improve spatial precision, for multielectrode TI strategy proposed here, this comes at the cost of a lower firing rate, which could be undesirable in many applications.

It is interesting to note that the fact spatial diversity hurts the precision of stimulation in TI, but can be harnessed to improve the precision in STIMULUS. This is especially evident when skull is more conductive (see Fig. 8). Significant difference of current density in different brain regions causes shallow neurons (near electrodes) to be engaged in TI, but STIMULUS is capable of creating precise stimulation patterns by harnessing this diversity. For lower skull-conductivity (*σ*_2_*/σ*_1_ = 1/80), the spatial diversity is less, and the accuracy of multielectrode TI is reasonable (engaging 174 voxels in total), although STIMULUS still outperforms multielectrode TI by stimulating only 124 voxels. But when skull has higher conductivity (*σ*_2_*/σ*_1_ = 1/15), spatial diversity deteriorates TI accuracy by stimulating 531 voxels, but STIMULUS can reduce engaged voxels to 126, a 4x improvement in accuracy in terms of volume stimulated.

We focused on required current densities for stimulation, but ignored the effect of higher current densities in the scalp (compared to the brain), e.g., on tissue health. Future work will incorporate constraints on maximum current densities and total currents.

We use idealized models, and the techniques need to be adapted to real head models. We expect this process to be similar to adapting EEG spherical head model techniques to real heads, and results on beamforming in real-head models [11] suggest that this should indeed be possible. This does require knowledge of conductivities in different parts of the head, which can be aided by obtaining a structural MRI scan. However, limited knowledge of conductivity of different layers will limit the accuracy of our techniques. Techniques that advance this understanding, e.g. electrical impedance tomography (EIT), improved imaging, and better estimation of tissue conductivity in vivo, can help with improved focusing. It is also possible that while capacitive effects are not prominent at 250 Hz (as reported in, e.g., [18]), they cause more significant phase shifts at higher frequencies such as thousands of hertz used in TI [1] and STIMULUS. However, this amounts to a small change in our optimization problem by incorporating phase changes induced by the medium at each location in addition to amplitude changes.

We note that the STIMULUS optimization is designed to provide a starting point to optimization approaches, and there are variations on the optimization formulation that could improve precision. E.g., instead of having foci for each patch at or very close to the target, as is done here, one may want all electrode-patches to focus slightly off-target, creating “virtual sources” at these foci that are closer to each other than the electrodes. One may also want a different frequency allocation across electrodes for both STIMULUS and multielectrode TI.

Further, we need to understand tradeoffs between firing rate and spatial precision. For multielec-trode TI, particularly when frequencies are allocated using our strategy, increasing spatial precision appears to be accompanied with reduced firing rate. However, the issue deserves a more thorough investigation. E.g., it is plausible that an increase in center frequency can allow frequency differences to be larger as well.

Finally, it is also possible to use STIMULUS with implanted electrodes for stimulating the peripheral nervous system (PNS), and also cochlear and retinal implants where localized stimulation is of immense importance. This needs to be explored deeply as well.

## 6 Acknowledgments

The authors acknowledge the support of the NSF CNS-1702694 and CMU BrainHUB. We also thank Alison Barth, Brent Doiron, Sanghamitra Dutta, Max Jin, Jana Kainerstorfer, Shawn Kelly, and Praveen Venkatesh for helpful discussions.

## Footnotes

1 This is analogous to dispersion of EEG signals going out of the brain through the skull and into the scalp [6, 7].

2 This assumption of all neurons having the same orientation, while seemingly stringent, yields pessimistic results in neural resolution because it disallows us from exploiting the diversity in orientation of different neurons to tailor firing [19].

3 It also depends on the radius of the head, but here that is assumed to be constant. The head-radius only scales all CID factors by the same amount.

